# Enhancing the efficacy, utility and throughput of the Transcription Block Survival peptide library screening platform

**DOI:** 10.1101/2025.03.28.645970

**Authors:** Andrew Brennan, T M Simon Tang, Jody M Mason

## Abstract

Genetically encoded peptide library screening is a powerful strategy for discovering inhibitors of protein-protein and protein-DNA interactions. The Transcription Block Survival (TBS) assay enables the *in vivo* selection of peptides that antagonise transcription factor (TF) binding by linking inhibition of DNA-binding to *E*.*coli* survival. However, previous TBS implementations required laborious re-engineering of the mDHFR coding region for each new target, limiting utility. Here, we present an enhanced and streamlined TBS platform that increases throughput, simplifies target switching and improves selection stringency. By relocating TF-DNA binding sites from within the mDHFR coding sequence into the mDHFR 5’-promoter/untranslated region, we preserve mDHFR folding and function, enabling rapid interchange of TF targets without the need for extensive construct redesign. We validated this system using three distinct TF targets, CREB1, ATF2 and DLX5, and two distinct consensus sites, demonstrating robust transcriptional block upon TF binding and efficient growth rescue upon peptide-mediated antagonism. Importantly, we expand the platform to accommodate full-length TFs, as exemplified by DLX5, allowing selection against biologically relevant multi-domain proteins. TBS continues to function exclusively by selecting for disruption of protein-DNA binding, ensuring mechanistic precision. Using this optimised TBS system, we successfully screened an 11.3-million-member peptide library to identify a potent antagonist of ATF2-CRE DNA binding within three months. This next generation TBS-platform dramatically improves screening efficiency and selection pressure while maintaining high biological relevance, providing a versatile and scalable tool for discovering functional peptide inhibitors of protein-DNA interactions with therapeutic potential.

## Introduction

Peptides represent a highly promising class of therapeutic molecules in drug discovery that can bridge the gap between small molecules and biologics, combining many favourable attributes.^1-5^ This unique positioning enables peptides to engage challenging drug targets that have been historically deemed undruggable. Advances in high-throughput genetically encoded peptide library screening technologies have further accelerated their discovery.^6-8^ These platforms operate by generating vast numbers of peptides which are either covalently linked or encapsulated with their corresponding nucleotide sequence. Peptide are then screened for their ability to bind or antagonise a target protein and hit sequences are rapidly identified by high-throughput sequencing.

The Transcription Block Survival (TBS) assay is a genetically encoded peptide library screening platform that selects for peptide antagonists of protein-DNA interactions in live bacterial cells.^7,9,10^ Unlike target binding selection assays such as mRNA display and phage display, a positive read-out from the TBS assay requires antagonism of target protein function (i.e. loss of DNA binding). TBS exploits the enzymatic activity of dihydrofolate reductase (DHFR) which is essential for bacterial DNA synthesis, and therefore cell growth. In TBS assay cells, endogenous *E. coli* DHFR activity is selectively inhibited by trimethoprim (TMP). Therefore, growth is dependent on expression of an exogenous mouse DHFR (mDHFR) gene from plasmid DNA, which is not inhibited by TMP. Introduction of target protein DNA binding sites into the coding region of the mDHFR and expression of the target protein, results in a steric transcriptional block of the mDHFR gene, and therefore abrogation of cell growth. Plasmid libraries encoding for highly diverse peptide sequences are transformed into TBS assay cells, meaning any cells producing peptides capable of disrupting the target protein-DNA interaction will remove the transcription block This restores expression of the mDHFR gene and confers a significant growth advantage. This operates as a molecular dial with peptide efficacy in antagonising target protein-DNA interaction correlating directly with bacterial growth rate, allowing competition selection.

Critically, peptide screening in a cellular context enables selection of other desirable attributes beyond target antagonism, including target specificity, solubility, tolerability, and biostability. Identifying peptides with desirable drug-like properties during the screening process reduces time and resources wasted on molecules with little or no subsequent therapeutic value. Moreover, millions of peptides can be tested in less than a litre of media using TBS, which is significantly more sustainable and cost effective than equivalent synthetic screening approaches. The TBS assay has produced effective inhibitors of oncogenic transcription factor (TF) cJun, which have shown efficacy in the reduction of cJun-driven melanoma cell viability.^10^ This demonstrated that efficacy observed during TBS can be translated into the mammalian cellular environment.

Although TBS library screening has been successfully employed to identify inhibitors of cJun,^7, 10^ BZLF1^9^ and other TFs (unpublished), a major limitation of the existing TBS screening protocol is the time and effort required to modify the reporter mDHFR gene for new targets that recognise and bind to different DNA sequences. As the target consensus sequence was introduced into the coding region of mDHFR, significant time and effort was required to design mutants with minimal changes (i.e. silent or conserved amino acids) to the protein sequence whilst introducing sufficient binding sites to produce the desired transcriptional block upon target protein expression. Further, our attempts to generate a TBS system for antagonism selection against cyclic AMP-responsive element (CRE)-binding TFs were unsuccessful. As we have done previously, a rational search was conducted to identify sites in the mDHFR gene which could be silently mutated to incorporate CRE sites, a second search was then conducted to identify sites which could be mutated to CRE where resulting substitutions were limited to conservative changes (i.e. similar sidechain properties) to surface exposed amino acids. This produced an mDHFR gene with five CRE sites in the coding region, but this was not sufficient to induce a significant transcriptional block and therefore the assay stringency was expected to be unacceptably low. Whilst an optimised construct could have been designed, the additional time investment required for target switching is undesirable for a broadly applicable screening platform.

To address this limitation, we report a streamlined and optimised strategy for rapid target switching in TBS. We validated this optimised TBS system using cyclic AMP-responsive element-binding protein 1 (CREB1),^11, 12^ activating transcription factor 2 (ATF2, also CREB2) ^13-15^ and distal-less homeobox 5 (DLX5)^16-18^ targets, demonstrating effective transcriptional block and growth restoration upon peptide antagonism. Further, we report on the utilisation of this enhanced TBS system to generate an effective peptide inhibitor of ATF2-DNA binding.

## Results and discussion

To enhance the efficacy and utility of TBS we have produced a modified assay design. Instead of mutating the translated region of the mDHFR gene, target DNA binding sites were instead introduced into the promoter and 5’ untranslated region (UTR) of the WT-mDHFR gene (**Figure 1**). Alteration of this region is facile, using standard cloning techniques to insert the desired sequence between *XhoI* and *SphI* restriction sites. As an initial investigation, two pES300d plasmids were constructed with either five or nine cAMP response element (CRE) consensus sites (**Figure S1, S2**). These constructs were tested to explore the number of CRE binding sites required to instigate a sufficient transcriptional block. Notably, CRE sites were introduced at the two lac operator sites in this plasmid. While mutations to the lac operator may impact repressor functionality, gene expression control by lac repressor/IPTG is not essential during TBS screening, since expression of the mDHFR reporter gene is controlled by the transcription block created by the target protein-DNA interaction. For the 5xCRE WT mDHFR construct a further three sites were added into the 5’ untranslated region. For the 9xCRE WT mDHFR construct, the length of the 5’ untranslated region was extended to accommodate six CRE sites, with an additional CRE site introduced in the 5’ region of the T5 promoter.

**Figure 1.**
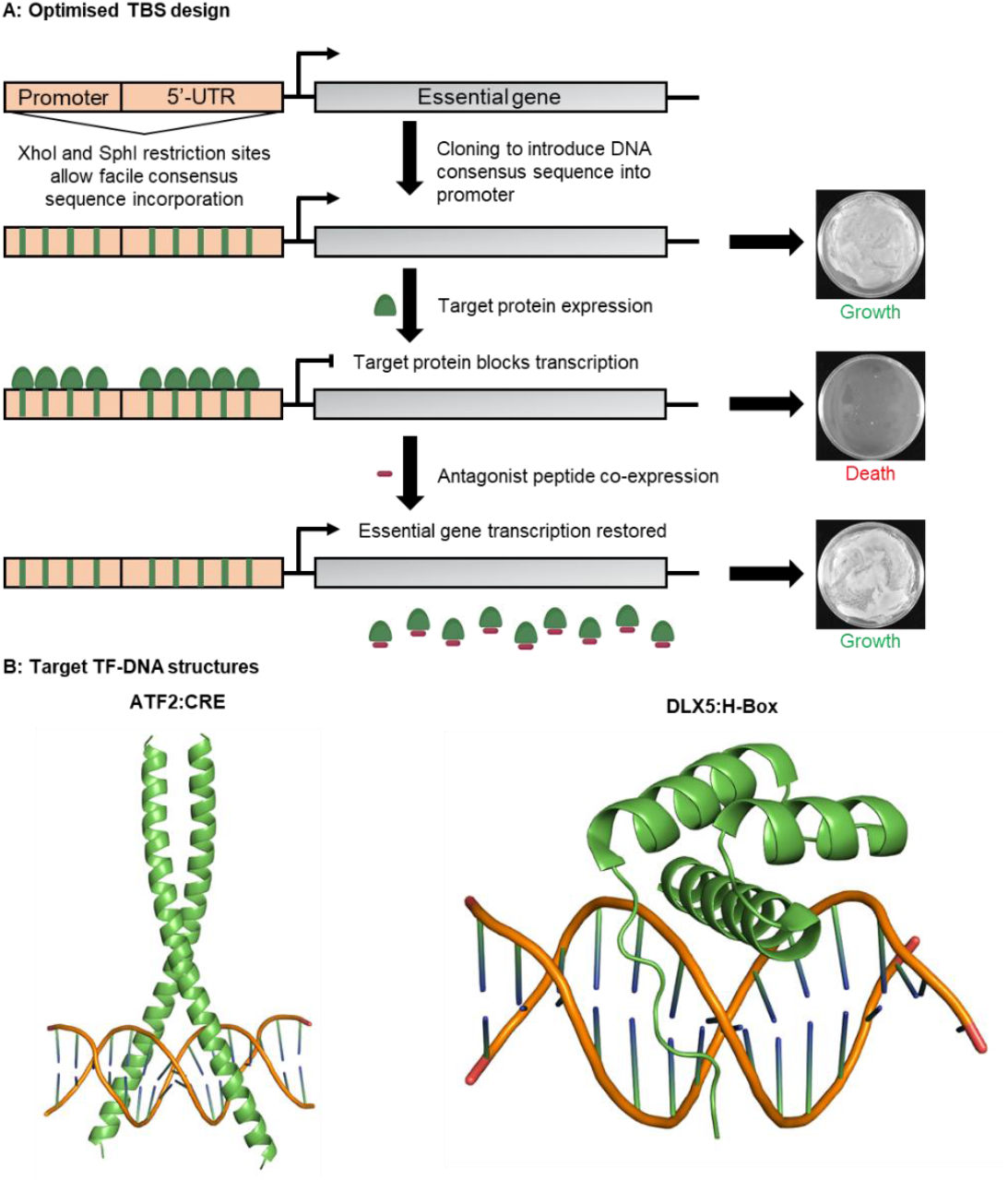
Streamlined TBS methodology facilitates rapid target switching with enhanced selection stringency against new oncogenic targets. (A) TBS assay design schematic illustrating the facile incorporation of target consensus sites into the 5’-promoter/UTR of the mDHFR gene. Upon target binding to these sites the essential gene transcription is blocked, abrogating bacterial growth. Introduction of an effective functional antagonist removes the transcription block and restores bacterial growth, allowing for a direct correlation between peptide antagonist efficacy and growth rate. (B) Alphafold prediction of the ATF2 bZIP domain homodimer bound to CRE DNA (pTM score: 0.85, iPTM score: 0.84, plDDT >90) and the crystal structure of the DLX5 homeodomain bound to H-Box DNA (PDB ID: 4RDU).

To validate the new TBS constructs, six BL21-Gold *E. coli* control strains were produced representing the theoretical steps required for the assay to operate, empty plasmids containing no gene of interest were employed to control for replication and antibiotic stresses (**Table 1**). Control strains 1 and 4 were transformed with the reporter plasmids containing the mDHFR gene, and two empty plasmids; control strains 2 and 5 were transformed with the reporter plasmids, target plasmid containing the CREB1 bZIP gene and an empty plasmid; control strains 3 and 6 were transformed with the reporter plasmids, target plasmid and the antagonist plasmid containing the A-CREB gene (peptide inhibitor of CREB1).^19^ Equal number of cells of each control strain were plated onto separate M9 agar plates supplemented with antibiotics (chloramphenicol, ampicillin, kanamycin), 1 mM IPTG and 8 µM TMP. Agar plates were incubated at 37°C for 24 hours before imaging.

**Table 1.** BL21-Gold E. coli control strains for TBS validation. Each control strain contains three pQE16 plasmid derivatives to facilitate reliable comparison of growth rates, with empty plasmids utilised where no expression is required.

| TBS Control Strain | pES300d | pES230d | pQE80 |
| --- | --- | --- | --- |
| 1 | 5xCRE mDHFR | Empty | Empty |
| 2 | 5xCRE mDHFR | CREB1 bZIP | Empty |
| 3 | 5xCRE mDHFR | CREB1 bZIP | A-CREB |
| 4 | 9xCRE mDHFR | Empty | Empty |
| 5 | 9xCRE mDHFR | CREB1 bZIP | Empty |
| 6 | 9xCRE mDHFR | CREB1 bZIP | A-CREB |
| 7 | 9xCRE mDHFR | ATF2 bZIP | Empty |
| 8 | 9xDLX mDHFR | Empty | Empty |
| 9 | 9xDLX mDHFR | DLX5 HD | Empty |
| 10 | 9xDLX mDHFR | DLX5 full length | Empty |

The significant growth of positive control strains 1 and 4 indicates that mDHFR was expressed and conferred resistance to TMP. In comparison, a significant drop in growth was observed for control strains 2 and 5, which were expressing the CREB1 bZIP gene. This confirmed the transcription block as a result of the protein-DNA interaction between CREB1 bZIP and CRE sites in the mDHFR gene. Finally, growth was restored upon expression of the A-CREB gene from the antagonist plasmid (strains 3 and 6). A-CREB is a known antagonist of the CREB1-CRE interaction^19^ which can remove the transcription block, enabling expression of mDHFR, therefore restoring TMP resistance and *E. coli* growth.

Strains transformed with the reporter plasmid containing 9xCRE sites showed a greater reduction in growth compared to those with 5xCRE sites (strain 5 *vs*. strain 2), indicating a correlation between the number of CRE sites and the degree of transcriptional shutdown. Since the enhanced transcription block generated greater stringency (i.e loss of cell viability) during TBS screening, subsequent investigations were performed using the reporter plasmid containing 9xCRE sites in the 5’-promoter/UTR region of the mDHFR gene. The same transcriptional block and growth inhibition were also observed when CREB1 was substituted with an alternative CRE-binding TF, ATF2 (**Table 1, strains 6 and 7; Figure 1B, 2**), demonstrating scope of application towards other protein targets that bind to CRE sites.

**Figure 2.**
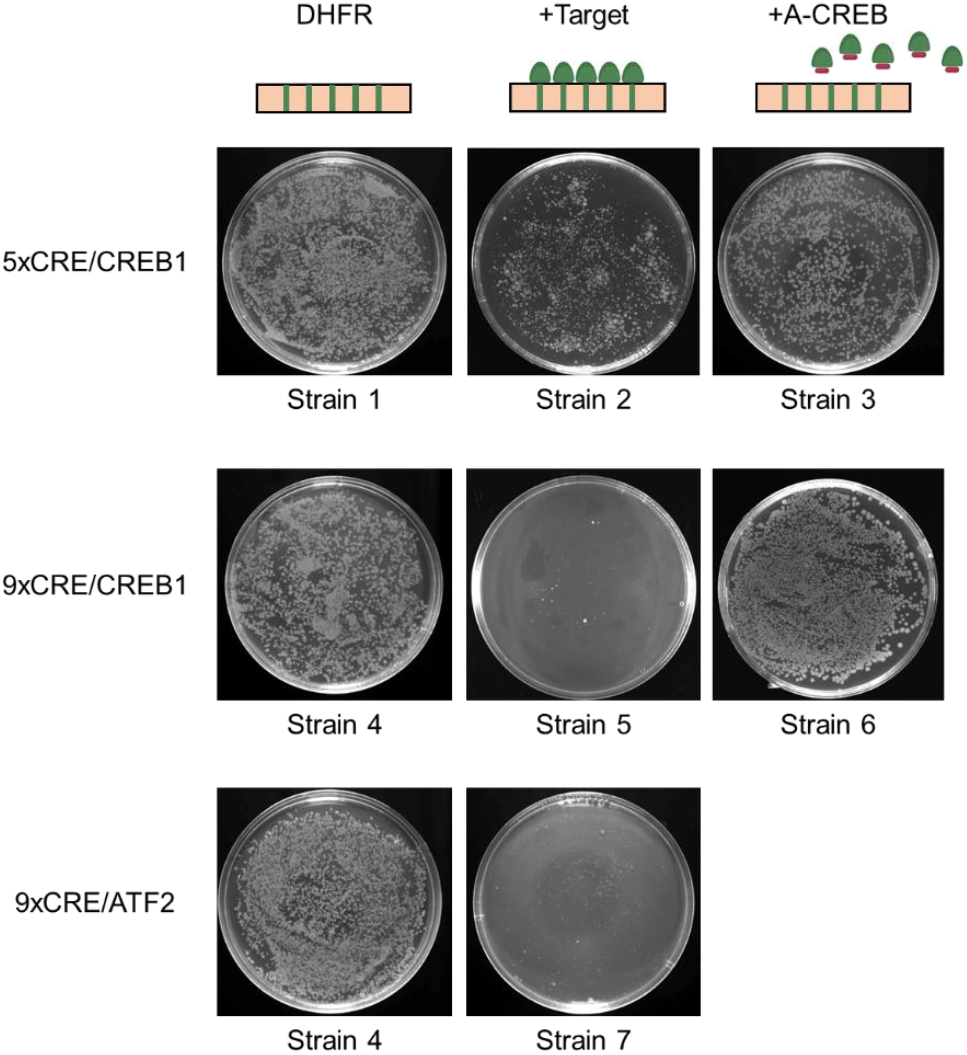
Introduction of CRE DNA sites into the 5’-promoter/UTR region of a mDHFR plasmid facilitates effective transcription block. Equal numbers of cells from the control strains (**Table 1**) were plated on selective M9 agar (8 µM TMP, 1 mM IPTG) and incubated before imaging.

To highlight the target switching capabilities for TF targets with different DNA recognition motifs, we next constructed a 9xhomeobox (H-Box) system for identifying inhibitors against the homeodomain-containing TF targets, such as the DLX (**Figure 1C**), pre-B cell leukaemia transcription factor (PBX) and orthodenticle homeobox protein (OTX) families. Mirroring the CRE system described above, 9xH-box consensus sequences were introduced into the 5’-promoter/UTR region of the mDHFR gene in the reporter plasmid (**Figure S3**).

The positive control, strain 8, transformed with the H-box containing reporter plasmid and empty target and antagonist plasmids was able to grow in TBS assay condition, which indicated expression of the mDHFR reporter gene to replace the TMP-inhibited endogenous DHFR activity (**Figure 3**). Consistent with the other TBS systems, introduction of the DLX5 homeodomain resulted in effective transcriptional block and subsequent reduction in the number of colony-forming units observed (strains 8 vs. 9).

**Figure 3.**
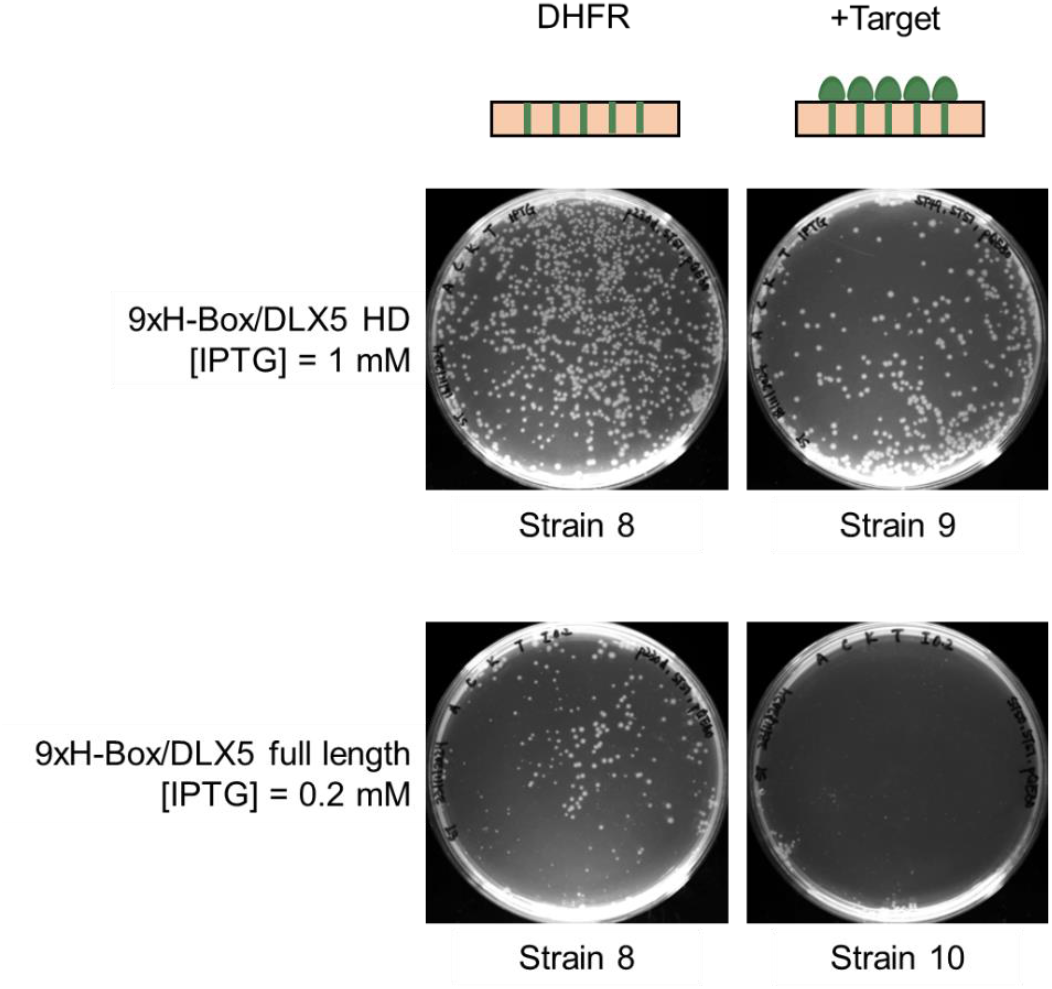
Introduction of H-box DNA sites into the 5’-promoter/UTR region of a mDHFR plasmid facilitates effective transcription block by both DLX5 full length and homeodomain. Equal numbers of cells from the control strains (**Table 1**) were plated on selective M9 agar (8 µM TMP, 1 or 0.2 mM IPTG) and incubated before imaging. Lower IPTG concentration was required for DLX5 full length due to overexpression-induced toxicity.

Many peptide library screening approaches require purified and immobilised target proteins, which can obscure potential binding sites, or be challenging for large intrinsically disordered proteins. Intracellular methods such as TBS overcome this limitation by enabling *in situ* peptide library screening during recombinant expression within *E. coli*, facilitating screening against any protein target without the need for purification, or the addition of fusions/tags. To demonstrate this advantage, the full length DLX5 gene was expressed from the target plasmid in the H-Box TBS system. However, overexpression of full length DLX5 protein demonstrated clear *E. coli* toxicity (**Figure S4** illustrates a difference in growth between strains 8 vs 10 upon 1 mM IPTG induction without TMP). To overcome this toxicity associated with overexpression of large, disordered proteins, IPTG concentration was reduced to 0.2 mM (**Figure S4** illustrates negligible difference in colonies at 0.2 mM IPTG). This reduction in IPTG reduced the number of colonies in selective conditions for Strain 8 (reporter plasmid only); however, a clear and significant reduction in bacterial growth, due to the transcriptional block, was also observed (**Figure 3**, Strain 8 *vs*. 10).

This validates the new TBS design for the introduction of a transcriptional block and for the ability of the positive control antagonist A-CREB to restore bacterial growth in the CRE/CREB systems. To demonstrate this new TBS designed retains its capacity for peptide library screening to generate effective TF-DNA antagonists, a 11.3 million member peptide library was generated for screening against ATF2, to *de novo* generate a peptide ATF2 antagonist for the first time. This library utilises an N-terminal acidic domain, originally developed by Vinson and co-workers, which is designed to interact with bZIP DBDs. This is followed by a semi-randomised LZ region. In this library the canonical coiled coil heptad repeat **e** and **g** positions were semi-randomised to include glu, gln or lys options. The two exceptions for increased library diversity were for: **e7** which were ile, asn, asp or val; and **g8** which were leu, gln, glu or val. The options at LZ **a** positions were leu, ile or val to allow for selection of improved hydrophobic packing in the coiled coil core, with **a6** the exception which had ile, asn, asp or val options to investigate potential core polar interactions. The **d** positions were not randomised as leu will favour formation of the desired parallel, dimeric coiled coil. Non-interfacial **b, c** and **f** positions in this region were fixed as a mixture of alanine to induce helicity, polar residues to produce an amphipathic water-soluble peptide and positively charged residues to enhance the cell-penetrance of the resulting peptide hit. Whole plasmid PCR library generation and subsequent transformation produced 55.5 million colony-forming units, with a 99.3% probability of full library coverage.

Upon transforming the library into BL21 Gold *E. coli* (already containing p300 9xCRE mDHFR and p230 ATF2 bZIP plasmids), a lawn of colonies was observed on the assay plates, with dilution plates indicating a 99.8% probability of full library coverage. After nine selection passages of continuous bacterial growth, a single assay winner peptide was observed, ATF2W (**Figure 4, S5**). ATF2 and ATF2W were synthesised by standard Fmoc-protected solid phase peptide synthesis and purified by RP-HPLC for biophysical characterisation (**Figure S6, S7**).

**Figure 4.**
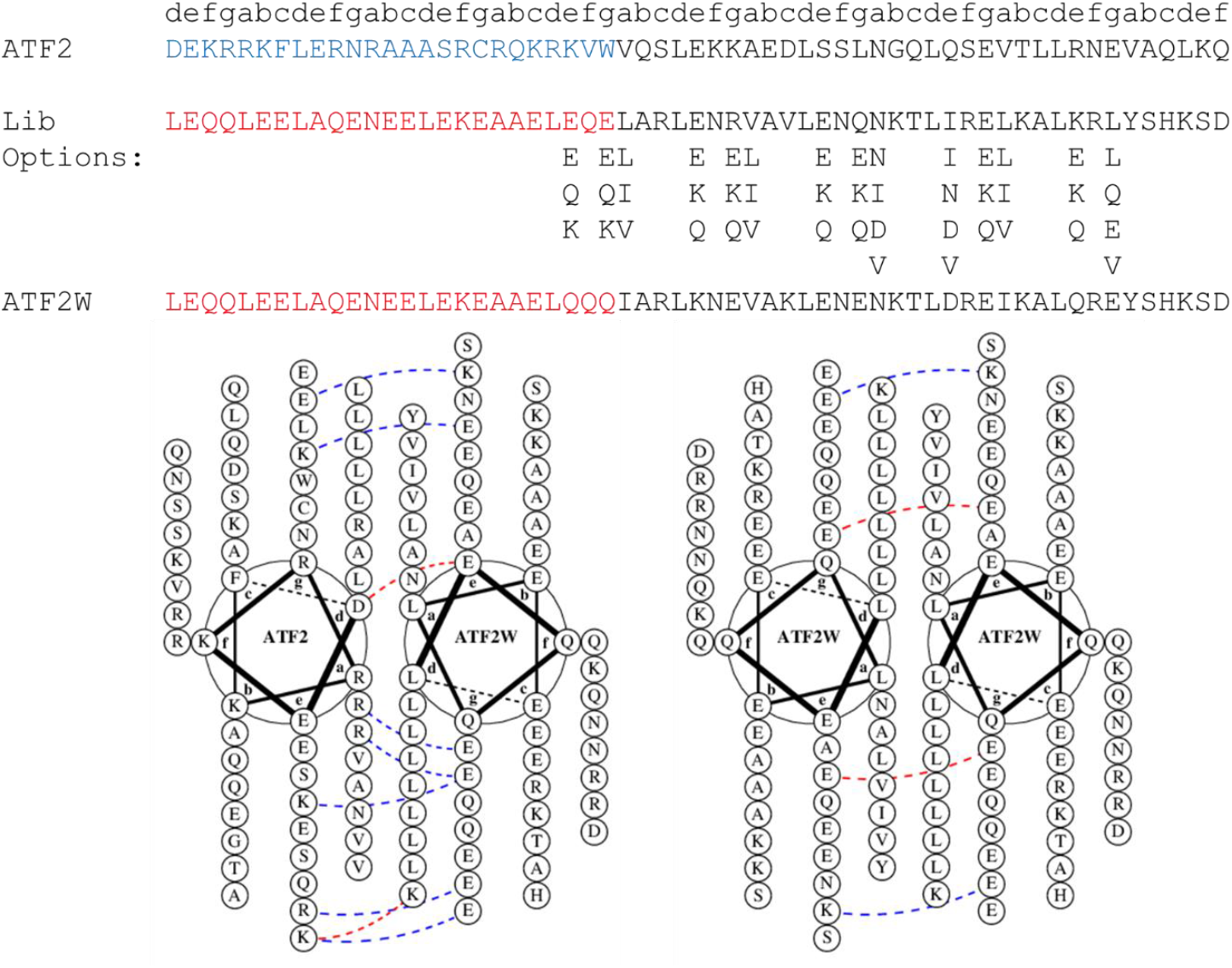
Target ATF2, antagonist library design and TBS assay winner peptide ATF2W sequences. Helical wheel diagrams illustrate the desired ATF2-ATF2W interaction and undesired ATF2W-ATF2W homodimer interaction.

**Figure 5.**
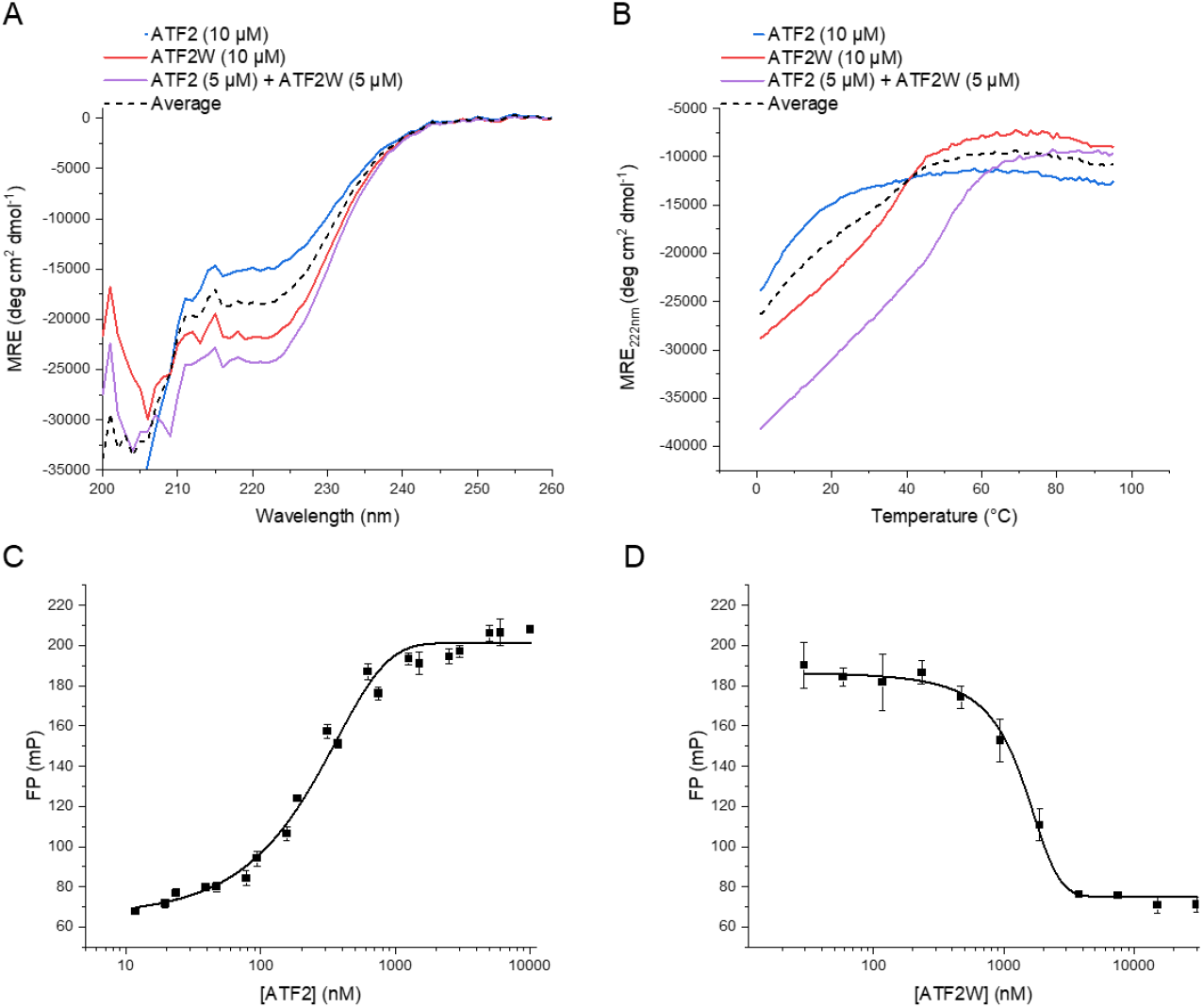
Biophysical characterisation of the ATF2-ATF2W interaction. (A) CD spectra of ATF2 and ATF2W indicating an increase in helicity upon mixture of the two components, indicative of the formation of a heterodimeric coiled coil. (B) CD thermal denaturation experiment indicates a T_m_ shift upon mixture of ATF2/ATF2W, supporting the formation of a stable heterodimer. The “average” CD trace in A and B represents an averaged value of the two unbound components which would be the trace for no interaction. (C) FP binding assay to illustrate ATF2 binding to FAM-CRE DNA with a K_D_=277±19 nM. (D) FP antagonism assay shows that ATF2W is capable of inhibiting the ATF2-CRE DNA interaction, at the top concentrations reducing the FP value to that of free DNA.

ATF2 and ATF2W binding was first characterised using circular dichroism spectroscopy, to measure changes in global secondary structure upon binding. CD spectra for both individual peptides (10 µM) and for a ATF2/ATF2W mixture (5 µM of each) were collected and converted to mean residual ellipticity. Total peptide concentration was constant in all samples as dimer helicity is concentration dependent. An average spectrum of the individual components was determined, as this represents the expected spectrum in the case of no interaction. All sample spectra displayed a variable minimum at 222 nm indicative of α-helicity. There was a significant 32% increase in helicity upon mixing of the two peptides (compared to the average), indicative of a coiled coil binding interaction. During these experiments, 10 mM TCEP was required as a reducing agent (1 hour incubation) to prevent ATF2 disulphide formation, which produced the significant noise in the spectra. To further investigate this binding interaction, a thermal denaturation experiment was performed on these samples, monitored the signal at 222 nm between 1-95 °C. The heterodimeric ATF2/ATF2W samples displayed a *T*_*m*_ of 55°C, a *ΔT*_*m*_ of 18°C relative to the ATF2W only sample. This significant increase in helicity and thermal stability indicates the formation of a helicity-inducing coiled coil binding interaction between ATF2 and ATF2W.

Fluorescence polarisation (FP) was next utilised to confirm that ATF2W binding produced the designed antagonism of the ATF2/CRE interaction. First, ATF2 (12 nM-10 µM) was incubated with FAM-CRE DNA (10 nM), and subsequent FP measurement indicated that ATF2 bound to CRE with a *K*_*D*_ = 277±19 nM. To measure ATF2W antagonism, an ATF2 concentration corresponding to 80% of the observed ΔFP was used (625 nM), incubated with CRE DNA (10 nM) and ATF2W (29 nM-30 µM) and the resulting FP measurement indicated that increasing ATF2W reduced the observed FP values, as DNA is released, with an *IC*_*50*_ = 1346±145 nM. Further, FP was utilised to confirm that ATF2W does not interact with CRE DNA (**Figure S8**).

## Conclusions

The TBS platform is a highly powerful intracellular peptide screening system for identifying peptide-based antagonists of TFs. It has been successfully employed to generate effective peptides that disrupt TF-DNA interactions, leading to effective inhibition of target function.^7,9,10^ However, earlier iterations of TBS required laborious redesign of the mDHFR reporter gene to incorporate new DNA recognition sequences for each TF, limiting the platform’s adaptability and broader application to other TF targets.

Our studies represent a significant advancement of the TBS screening platform by overcoming previous limitations through strategic redesign of the reporter plasmid to enable rapid and flexible target switching. In this improved system, TF binding sites were relocated from the translated region of the mDHFR gene into the 5’ UTR and promoter region. This means that the 5’ UTR and promoter region is the only region that requires editing ahead of TBS screening. This design eliminates the need to re-design the coding sequence each time where changes could compromise reporter function and assay integrity. Additionally, the use of WT-mDHFR in place of the previously modified TRE-mDHFR variant results in increased enzymatic activity, promoting faster bacterial growth and thereby accelerating the TBS screening process.

The current study investigated the use of five or nine DNA recognition sites to incur the required transcription block in a CREB1/CRE DNA system. Reporter plasmids containing nine binding sites in the 5’ UTR and promoter were proven to generate highly effective transcription blocks for both CRE- and H-Box mDHFR. Further optimisation of number and placement of DNA binding sites could be performed to balance growth rate and stringency during selection. CREB1 and ATF2 were employed to evaluate the utility of the CRE-mDHFR reporter plasmid against different CRE-binding TFs. Reduction in *E. coli* growth, indicating effective transcription blocks were observed in both instances. As a proof-of-concept, an 11.3-million-member peptide library was screened against ATF2 by TBS using the CRE-mDHFR reporter plasmid. This resulted in the selection of ATF2W, capable of binding to ATF2 and sequestering it from its cognate CRE DNA. Future work will utilise these systems to generate CREB1 and DLX5 antagonists.

A reporter plasmid containing nine H-box DNA sequences in the 5’ UTR and promoter region was created to demonstrate adaptability of the new reporter plasmid design, enabling screening against homeodomain-containing TFs. Expression of DLX5 genes encoding for the homeodomain or the full-length TF both resulted in the expected reduction in E. coli growth. Notably, this exemplar highlights the potential applications of the TBS screening platform against a different class of DNA-binding domains.

In summary, the optimised TBS platform, enabled by a re-designed reporter system, allows for rapid and efficient target switching, and was successfully applied to three TFs across two distinct cognate DNA recognition sequences. The updated plasmid design not only simplified adaptation to new targets but also enhanced selection pressure and increased growth rates during screening, significantly improving throughput. Collectively, these improvements enabled the identification of an effective antagonist of ATF2-CRE DNA binding within just three months (from cloning and TBS validation, through library construction and screening, to peptide synthesis and biophysical characterisation), demonstrating the platform’s powerful utility for intracellular peptide discovery.

## Acknowledgments

JMM is grateful to the Medical Research Council (MR/T028254/1) and the Biotechnology and Biological Sciences Research Council (BB/X001849/1, and BB/T018275/1).

