## Supplementary Information for "Enhancing the efficacy, utility and throughput of the Transcription Block Survival peptide library screening platform"

### Experimental

**Plasmid constructs**: Target cognate binding sequence-containing inserts were produced using PCR fill-in reactions from synthesised primers (Merck). They were then subcloned using *XhoI* and *SphI* sites into the pQE16-derivative pES300d WT mDHFR plasmid (**Figure S1, S2, S3**). Target constructs were subcloned into pQE16 derivative plasmid pES230d. The human ATF2 bZIP domain spans from Asp$^{352}$ to Gln$^{410}$, the CREB1 bZIP domain spans from Ala$^{283}$ to Asp$^{341}$, the DLX5 HD spans from Val$^{137}$ to Asn$^{198}$. A-CREB has the following sequence: LEQQLEELAQENEELEKEAEELEQELAELENRVAVLENQNKTLIEELKALKDLYCHKSD, and was subcloned into pQE16 derivative plasmid pQE80.

**TBS validation:** For validation experiments, overnight cultures of control strains were diluted and an equal number of cells were plated on M9 minimal media plates containing ampicillin, kanamycin and chloramphenicol to maintain the required plasmids. Four plate types were used: antibiotics only, antibiotics+16 µM TMP, antibiotics+0.2/1 mM IPTG, antibiotics+16 µM TMP+0.2/1 mM IPTG. These plates were used to investigate viability of the strains and confirm they grew equally, the overexpressed protein/peptides did not induce significantly different toxicity, that TMP inhibited bacterial growth and to apply selection pressure respectively.

**Library construction and screening:** Two primers containing degenerate base positions were purchased (Merck): ATF2L_F: GCTAAGCTTVARAACVARRWTAAGACCCTGRWTCGTVARVTYAAGGCGCTGVARCGTSWGTACAGCCACAAAAGCGATTAATGATAAAATTAGCTGAGCTTGGACTC; ATF2L_R: GTGAAGCTTAGCRABYTBGTTYTBTAAACGTGCRABYTBTTGYTBCAACTCGGCCGCTTCTTTC. Underlined sequences indicate degenerate codons. A whole plasmid PCR reaction was performed using these plasmids and the A-CREB pQE80 plasmid to generate the linear library plasmid. The two ends of the purified PCR product were digested with *HindIII* (NEB) and purified. The plasmid was cyclised using T4 ligase (Thermo) and purified before electroporation into TG1 cells (Agilent). Dilution plates were utilised to determine transformation efficiency and colonies were scraped and the library DNA was purified. The following equation was utilised to determine library coverage by the number of single colonies: $E = 100 \times (1 - \frac{1}{n})^m$ where E is the percentage of the library missing, m is the number of colonies collected and n is the library size. Library DNA quality was assessed by sequencing both the DNA pool and a number of single colonies to show degenerate codons in the correct positions in the pool and to show a diversity of library members from single colonies.

The pool of library DNA was transformed into BL21-Gold cells already containing pES300d-9xCRE mDHFR and pES230d-ATF2 bZIP. The library transformants were first plated out onto selective agar plates (M9 media, ampicillin, kanamycin, chloramphenicol, TMP, IPTG) and grown at 37°C for 16h. Colonies from this first round of selection were pooled and serially grown in liquid culture at starting $OD_{600}$ of 0.05 and grown at 37°C with shaking at 200 rpm until the $OD_{600}$ reached 0.6. At each passage step, a sample of the culture was plated on LB agar (supplemented with antibiotics to maintain plasmids) to select and sequence individual colonies, and a DNA pool was also sequenced. This allows the occurrence of library members to be monitored as winner sequences are selected for.

**Peptide synthesis**: Peptides were synthesised using a Liberty Prime microwave peptide synthesiser (CEM) at a 0.1 mmol scale on rink amide ProTide (low loading) resin using standard Fmoc solid-phase methodology. Coupling was performed in a 5 mL reaction using 5x amino acid, 5x Oxyma Pure and 10x N,N'-diisopropylcarbodiimide in dimethylformamide (DMF). Deprotection was performed by addition of 0.75 mL 30% pyrrolidine into the reaction. Peptides were acetylated at the N-terminus by a final reaction with 3x acetic anhydride, 4.5x diisopropylethylamine in DMF for one hour at room temperature. Incubation in a cleavage mixture (95% TFA, 2.5% triisopropylsilane, 2.5% H2O, 10 mL) for 4 h at room temperature cleaved the peptide from the resin and removed side chain protecting groups. 2.5% DODT was added to the cleavage cocktail for ATF2. The resin was removed by filtration and cleaved peptides were precipitated in diethyl ether at -80°C and centrifuged. This pellet was washed a further four times with diethyl ether before it was dried overnight at room temperature. Peptides were resuspended in 1:1 water:acetonitrile (0.1% TFA) before purification using RP-HPLC with a Jupiter Proteo column (4-μm particle size, 90 Å pore size, 250 × 10 mm; Phenomenex) using a water:acetonitrile gradient (0.1% TFA). Peptide masses and purity (>95%) were verified by electrospray ionisation mass spectrometry. Upon solubilising in buffer, peptides were centrifuged to ensure insoluble/aggregated material was removed before peptide concentration determination.

**Circular Dichroism:** An Applied Photophysics Chirascan was used for CD measurements, with a 200 µL sample in a 1 mm path length CD cell. Peptide samples were suspended in 20 mM potassium phosphate,150 mM potassium fluoride, 10 mM TCEP.HCl at pH 7.4. Three scans between 190 and 260 nm were collected with a bandwidth of 1 nm and data sampled at a rate of 0.5 $s^{-1}$. These scans were averaged and converted to molar residue ellipticities (MRE). Thermal denaturation experiments were performed by measuring the ellipticity at 222 nm over a 1 to 95°C gradient at 1°C increments. Post-melt scans at 20°C confirmed the transitions were reversible as they overlaid within 5% of the pre-melt scan. The resulting thermal denaturation curves were converted to MRE and fitted to a two-state model, derived via modification of the Gibbs–Helmholtz equation to determine the melting temperature ($T_m$).

**Fluorescence polarisation**: Fluorescence polarisation was measured in a CLARIOstar fluorescence microplate reader (BMG Labtech), using black polystyrene, non-binding surface 96-well half area plates (Corning) in a 20 mM potassium phosphate, 50 mM sodium chloride, 5 mM MgCl2, 10 mM TCEP, pH 7.4 buffer. Palindromic FAM-CRE DNA oligo was purchased from Merck: 5'[6FAM]- GCGCTGACGTCAGCGC. 200 µM of the oligo was prepared in water and then annealed by heating to 95°C for 10 minutes before cooling slowly to room temperature to give a 100 µM duplex stock. Samples containing 10 nM FAM-CRE and a serial dilution of ATF2 (12 nM-10 µM) were prepared and incubated at room temperature for 1 hour before transferring to the assay plate for measurement. Data were fit to a one site binding model. For antagonism experiments, the ATF2 concentration required for 80% response was selected. Samples containing 10 nM FAM-TRE, 625 nM ATF2 and a serial dilution of ATF2W (29 nM-30 µM) were prepared and incubated at room temperature for 2 hours before transferring to the assay plate for measurement.

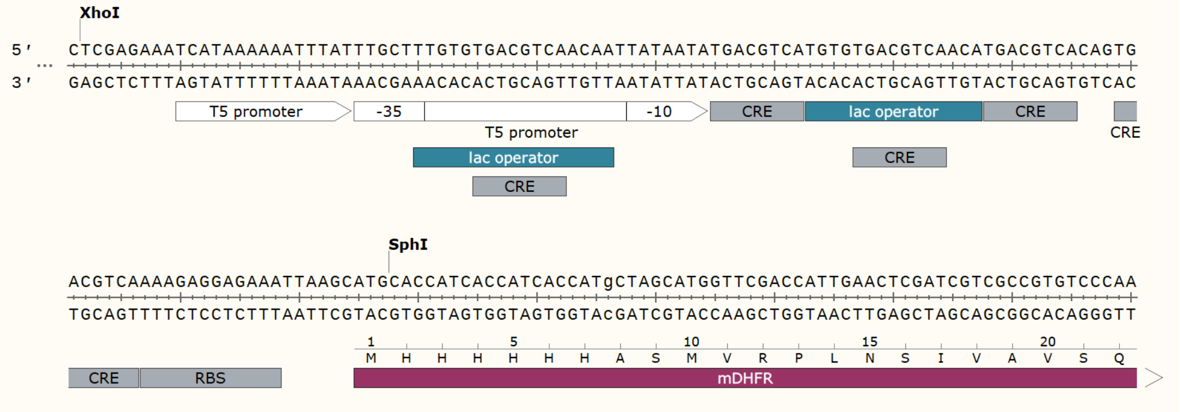


**Figure S1** - 5’-promoter/UTR DNA sequence of the 5xCRE mDHFR pES300d plasmid, indicating the insertion of CRE sites to facilitate transcriptional block.

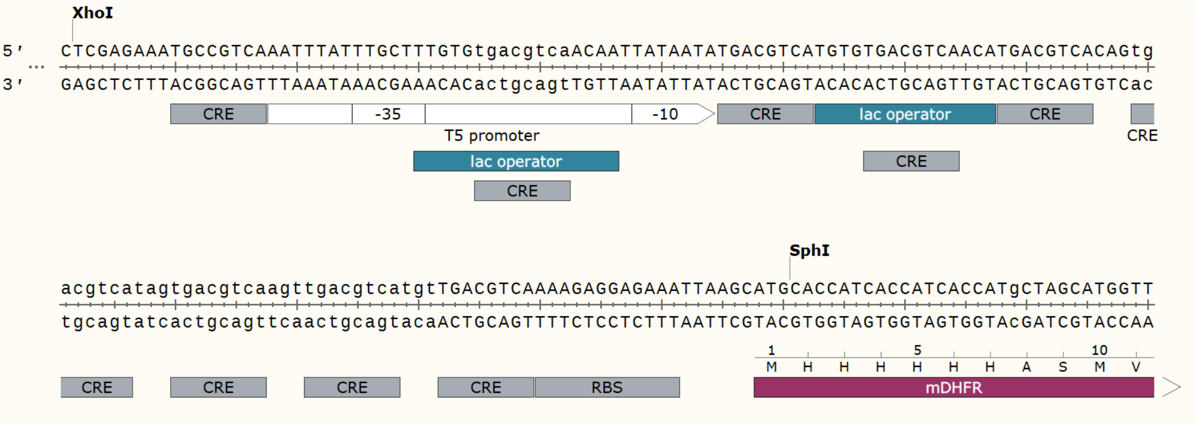


**Figure S2** - 5’-promoter/UTR DNA sequence of the 9xCRE mDHFR pES300d plasmid, indicating the insertion of CRE sites to facilitate transcriptional block.

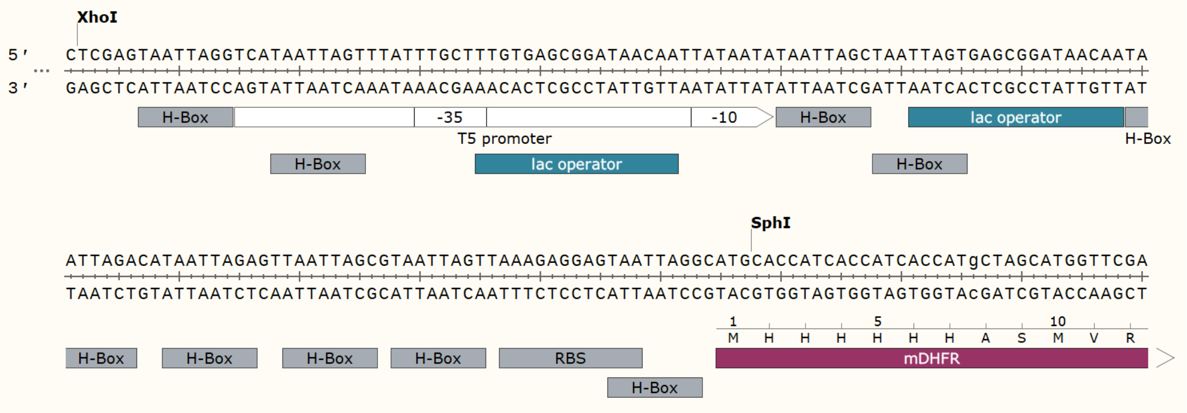


**Figure S3** - 5’-promoter/UTR DNA sequence of the 9xH-Box mDHFR pES300d plasmid, indicating the insertion of H-Box sites to facilitate transcriptional block.

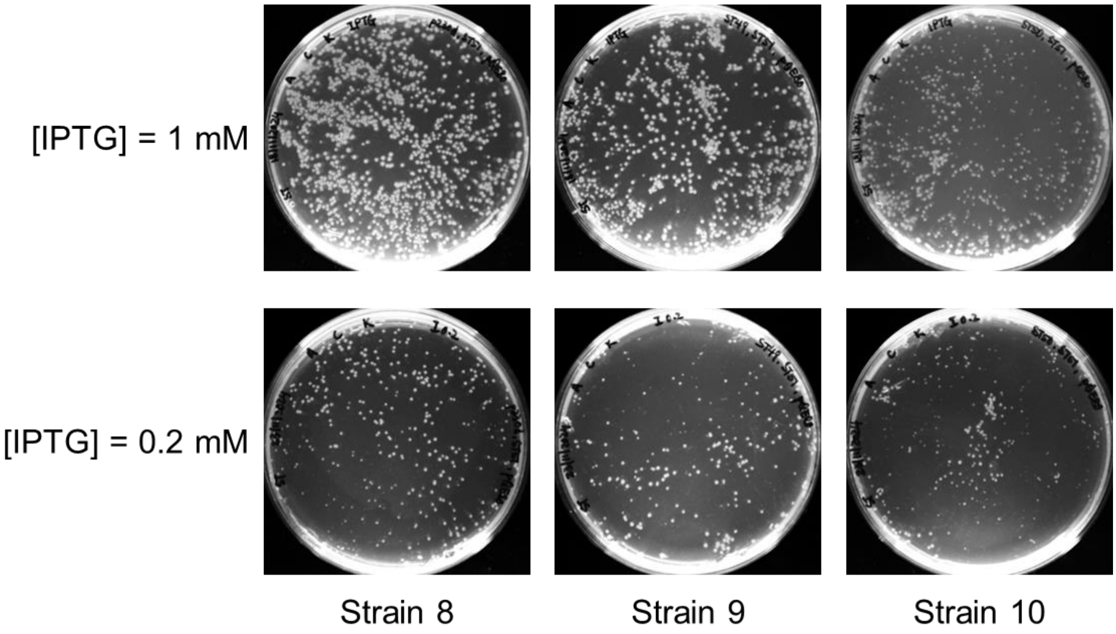


**Figure S4** - Equal numbers of cells from control strains 8-10 were plated on M9 agar + 0.2 or 1 mM IPTG (no TMP). This illustrated a reduction in colonies for strain 8/9 vs 10 at 1 mM IPTG due to the toxicity associated with overexpression of the large and disordered full length DLX5 in strain 10. Reducing IPTG concentration to 0.2 mM produced no significant difference in control strain growth, due to the decreased toxicity burden of protein overexpression.

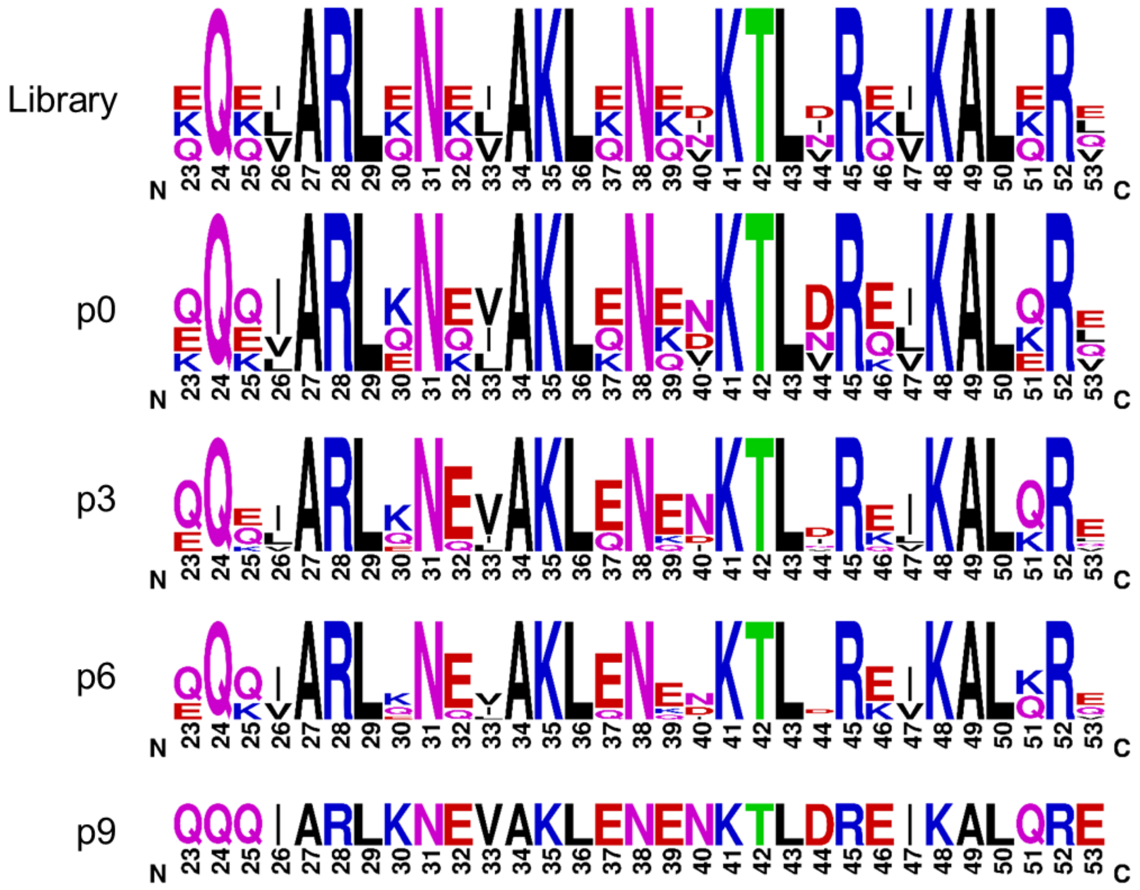


**Figure S5 -** Sequence logos showing the relative abundances of the amino acid library options at various passage numbers during screening, determined from individual colony sequencing. Small sample sizes may not be fully representative of the library diversity but simply indicate sequences which were observed.

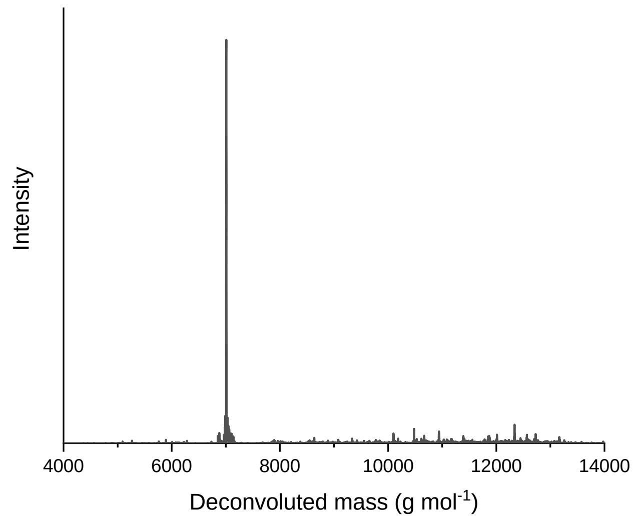

**Figure S6** - MS of purified ATF2 bZIP indicating a MW of 7005.62 g mol$^{-1}$ (predicted: 7004.82 g mol$^{-1}$).

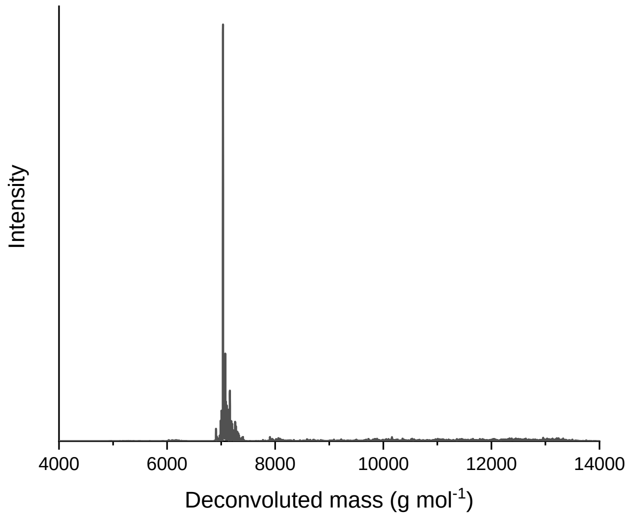


**Figure S7** - MS of purified ATF2W indicating a MW of 7031.42 g mol$^{-1}$ (predicted: 7029.59 g mol$^{-1}$).

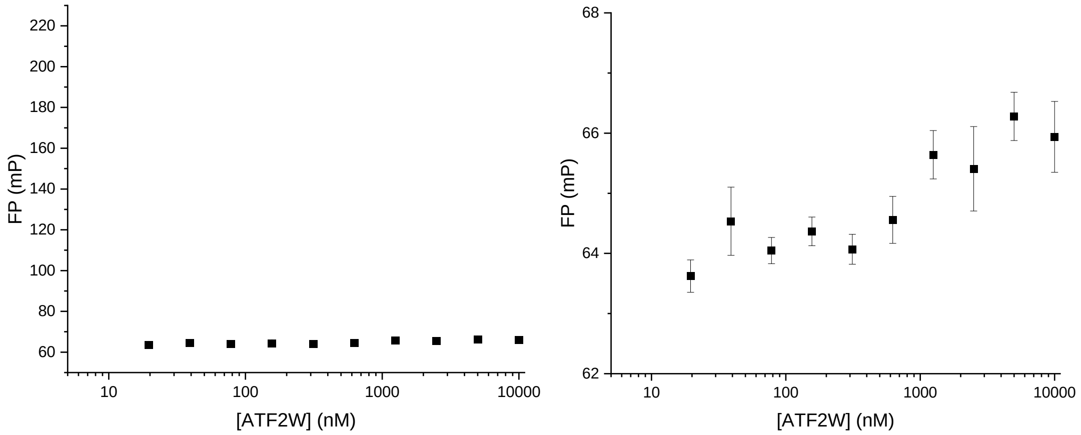


**Figure S8** - FP binding assay illustrates that ATF2W (9 nM-10 μM) does not bind to FAM-CRE DNA (10 nM), up to a 1000x excess. Plotted on the same scale as **Figure 5C** (left) and at a reduced Y-axis range (right).
